# Large-Scale Structure and Individual Fingerprints of Locally Coupled Sleep Oscillations

**DOI:** 10.1101/289157

**Authors:** Roy Cox, Dimitris S Mylonas, Dara S Manoach, Robert Stickgold

## Abstract

Slow oscillations and sleep spindles, the canonical electrophysiological oscillations of non-rapid eye movement sleep, are thought to gate incoming sensory information, underlie processes of sleep-dependent memory consolidation, and are altered in various neuropsychiatric disorders. Accumulating evidence of the predominantly local expression of these individual oscillatory rhythms suggests that their cross-frequency interactions may have a similar local component. However, it is unclear whether locally coordinated sleep oscillations exist across the cortex, and whether and how these dynamics differ between fast and slow spindles, and sleep stages. Moreover, substantial individual variability in the expression of both spindles and slow oscillations raise the possibility that their temporal organization shows similar individual differences. Using two nights of multi-channel electroencephalography recordings from 24 healthy individuals, we characterized the topography of slow oscillation-spindle coupling. We found that while slow oscillations are highly restricted in spatial extent, the phase of the local slow oscillation modulates local spindle activity at virtually every cortical site. However, coupling dynamics varied with spindle class, sleep stage, and cortical region. Moreover, the slow oscillation phase at which spindles were maximally expressed differed markedly across individuals while remaining stable across nights. These findings both add an important spatial aspect to our understanding of the temporal coupling of sleep oscillations and demonstrate the heterogeneity of coupling dynamics, which must be taken into account when formulating mechanistic accounts of sleep-related memory processing.

## Introduction

During non-rapid eye movement (NREM) sleep, highly organized oscillatory rhythms of slow oscillations and sleep spindles occur across widespread brain areas. In recent years, these electroencephalographic (EEG) waveforms have attracted considerable attention, owing to their close relation with cognitive functioning ^1,2^ Both spindles and slow oscillations (SOs) are closely linked to sensory information gating ^3–6^, and plasticity and memory processes ^7–11^. Moreover, their altered expression in aging and various neuropsychiatric disorders ^12–17^ has made them therapeutic targets of considerable interest.

SOs are large-amplitude ~1 Hz neocortical oscillations of alternating depolarized and hyperpolarized brain states that modulate neuronal spiking ^18–20^. Conversely, spindles are short (0.5–2 s) bursts of sigma band activity (9–16 Hz) initiated by the thalamus that propagate to cortex ^21^. In humans, spindles can be classified as either slow (9–12.5 Hz) or fast (12.5–16 Hz; but see ^22^ for an overview of spectral definitions in use). Aside from their oscillatory frequency, fast and slow spindles differ in topography ^23,24^, hemodynamic activity ^25^, heritability ^26^, and development ^27^. Moreover, spindles exhibit replicable individual differences in frequency ^26,28^ and topography ^22^, relating to underlying variability in neuroanatomy ^29^.

Intriguingly, SOs and spindles are temporally coordinated, such that spindle activity preferentially occurs in a particular phase of the SO cycle. Scalp recordings have repeatedly demonstrated that fast spindles tend to have maximal power around the depolarized SO peak, while slow spindles have greatest intensity in the peak-to-trough transition ^30–33^. Intracranial recordings revealed similar SO-coupling dynamics for fast spindles across many neocortical regions ^34^, as well as in hippocampus ^35^ and thalamus ^36^. This phenomenon of cross-frequency coupling ^37^ has been tied to sleep-dependent memory consolidation, such that appropriate spindle timing relative to the SO phase enhances memory ^16,38–40^.

While both SOs and spindles can be observed across the cortex, they are more often local (i.e., restricted in spatial extent) than global events ^20,41–46^, potentially allowing for circuit-specific plasticity and consolidation processes ^33,47–49^. Indeed, locally detected EEG SOs coordinate spindles in a spatially restricted fashion ^31^. However, the spatial extent of local SO-spindle coupling and its compatibility with regionally specific memory processing remain unknown. Moreover, assessing coupling dynamics separately for fast and slow spindles, and for deep N3 and light N2 NREM sleep, could shed light on the functional role of these different spindle classes and sleep stages. Finally, observations of reproducible individual differences in the expression of spindles, as well as SOs ^45^, raise the possibility that SO-spindle coupling dynamics show similar individual differences.

Here, we demonstrate that scalp SOs are spatially restricted, yet coordinate local spindle activity at virtually every cortical site. However, important variations can be seen depending on spindle class, sleep stage, and cortical region. Moreover, we show the presence of marked individual differences in the SO phase at which spindle activity is maximal. Although this variability was not associated with overnight procedural memory improvement, it was highly stable across nights, indicating that coupling phase constitutes a stable trait that may have important functional and clinical implications.

## Methods

### Protocol and participants

The present study describes novel analyses of full-night EEG data that we reported on previously ^22^. Twenty-four healthy individuals (age: 30.2 ± 6.3; 18 male, 6 female) gave written informed consent in accordance with the Declaration of Helsinki and were paid for participation. Data were acquired as part of a double-blind, placebo-controlled, cross-over study of eszopiclone in schizophrenia patients. Only the two consecutive placebo nights of the control group are considered in the present study. The study was approved by the Partners Human Research Committee. Additional details regarding participant screening and the protocol can be found in our previous report ^22^.

### Finger tapping Motor Sequence Task

After an initial baseline night, on the second night participants performed the finger tapping Motor Sequence Task (MST), a well-validated probe of sleep-dependent memory consolidation ^38,50–53^. Subjects were trained on the MST 2 h 45 m prior to bedtime and tested 1 h after awakening. The MST involves pressing four numerically labeled keys on a standard computer keyboard with the fingers of the left hand, repeating a five-element sequence (4-1-3-2-4) "as quickly and accurately as possible" for 30 seconds. The numeric sequence was displayed at the top of the screen, and dots appeared beneath it with each keystroke. During both training and test sessions, participants alternated tapping and resting for 30 seconds for a total of 12 tapping trials. The outcome measure was the number of correct sequences per trial, which reflects both the speed and accuracy of performance. Overnight improvement was calculated as the percent increase in correct sequences from the last three training trials to the first three test trials the following morning ^52^.

### Data acquisition and preprocessing

Polysomnography was collected using 62-channel EEG caps (Easycap GmbH, Herrsching, Germany) with channel positions in accordance with the 10–20 system. Additional EEG electrodes were placed on the mastoid processes, and on the forehead as online reference. Electrooculography (EOG) and bipolar chin electromyography (EMG) were monitored as well. An AURA-LTM64 amplifier and TWin software were used for data acquisition (Grass Technologies). Impedances were kept below 25 kΩ and data were sampled at 400 Hz with hardware high-pass and low-pass filters at 0.1 and 133 Hz, respectively. Sleep staging was performed in TWin using 6 EEG channels (F3, F4, C3, C4, O1, O2) referenced to the contralateral mastoid (bandpass filtered 0.3–35 Hz), bipolar EOG (0.3–35 Hz) and bipolar EMG (10–100 Hz), on 30 s epochs according to AASM criteria ^54^.

Initial processing of multi-channel EEG data was performed in BrainVision Analyzer 2.0 (BrainProducts, Germany). All EEG channels were band-pass filtered between 0.3 and 35 Hz and notch filtered at 60 Hz. Channels displaying significant artifacts for more than 30 minutes of the recording were interpolated with spherical splines. EEG data were then re-referenced to the average of all EEG channels. Upon visual inspection, epochs containing artifacts were removed. To remove cardiac artifacts we used independent component analysis with the Infomax algorithm ^55^. For each night and individual, remaining epochs were concatenated separately for sleep stages N3 and N2, resulting in 80.2 ± 39.5 (mean ± SD) and 82.3 ± 28.7 min of available N3 for the two nights, and 174.4 ± 60.1 and 211.4 ± 52.4 min of N2.

All subsequent processing and analysis steps were performed in Matlab (the Mathworks, Natick, MA), using custom routines and several freely available toolboxes including EEGlab ^56^ and the CircStat toolbox for circular statistics ^57^. After removal of non-EEG channels and the mastoids, leaving 58 channels for analysis, we applied a surface Laplacian filter to each record to both minimize the impact of volume conduction and thereby highlight local cortical processing ^58,59^, and allow more accurate estimates of oscillatory phase ^60^.

### SO detection

Individual SOs were detected during N3 and N2 on each channel using an established method closely resembling our previous approach ^31^. Specifically, the Laplacian-filtered signal of each channel was band-pass filtered between 0.4 and 1.5 Hz (zero-phase shift, third-order IIR filter). An SO was detected when (1) the latency between subsequent negative and positive zero-crossings of the filtered trace fell between 0.3 and 0.75 s, (2) the negative half-wave reached a minimum of −1 μV/cm^2^, (3) the amplitude difference between the trough and the subsequent local maximum exceeded 2 μV/cm^2^, and (4) the unfiltered EEG amplitude difference between the time points corresponding to the Laplacian-based SO trough and peak exceeded 50 μV. This final criterion ensures that Laplacian-detected SOs correspond to similar fluctuations in the regular EEG. The only difference from our previous report concerns the precise Laplacian and EEG amplitude criteria, which were relaxed here to detect events in regions where SO-band amplitude fluctuations are of smaller amplitude than in frontal areas, where SOs are conventionally detected. We marked SOs as 2 s time windows centered on each trough (1 s on either side) to capture at least one full cycle of the oscillation. Note that while this procedure allows for overlapping time windows between closely spaced SOs, the number of detected SOs per minute (see Results) suggests this occurred infrequently.

### Time-frequency power

To assess how local SOs modulate faster activity across an extended frequency range, we performed time-frequency analyses centered on each channel's SO troughs. Window size was set to ± 1.5 s around each SO trough to avoid edge artifacts stemming from the convolution. Decomposition was performed with a family of complex Morlet wavelets, according to 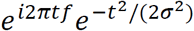, where *i* is the imaginary operator, *t* is time, *f* is frequency (30 logarithmically spaced frequencies between 5 and 25 Hz), and *σ* is the wavelet width. We defined the width *σ* as *λ*/(2*πf*/), where *λ* is the number of wavelet cycles, which was set to 5. The resulting time-frequency representations were down-sampled to 100 Hz to reduce the amount of data. Power was defined as the squared magnitude of the convolution result. Power estimates at each time point in each SO window were converted to percentage change relative to average power from −1 to 1 s surrounding the SO trough (i.e., single-trial baseline). This normalization ensures that power values can be compared across frequencies and channels. For single-subject analyses, time-frequency spectrograms were averaged across all SO windows for each channel. For group analyses, we further averaged across subjects.

### SO co-occurrences

We determined the degree to which SOs detected on a "source channel" were also observed on a "target channel". Specifically, for each subject, source channel, and SO, a co-occurrence was counted when a target channel (excluding the source channel itself) expressed an SO trough within a fixed window of either ±400 or ±100 ms surrounding the source SO trough. Target channel counts were averaged across SOs to obtain a measure of average channel involvement for each source channel. In a complementary approach, we determined, for each subject and source channel, the relative proportion of source SOs that were detected at each possible number of target channels (ranging from 1 to 57). These normalized histograms were subsequently averaged over subjects, and the resulting group-level histograms were converted to cumulative distributions to determine the maximum number of target channels involved for 50, 75, and 99% of source SOs.

### SO-spindle coupling

We obtained estimates of both SO-spindle coupling strength and coupling phase (defined below) for each subject, night, sleep stage, spindle class, and electrode. We first filtered the Laplacian-transformed multi-channel data in the canonical SO range (0.5–2 Hz), and in 1.3 Hz wide windows centered on each subject's individual fast and slow sigma frequencies. Individualized sigma frequencies were determined using a spatial filtering approach based on generalized eigendecomposition, as described in our previous report ^22^. We used the Hilbert transform to obtain the analytic signal of each channel in each frequency band, and extracted the instantaneous phase of the SO signals and the instantaneous amplitudes of the fast and slow sigma signals at every time point. Instantaneous amplitudes were squared to obtain power envelopes. Then, for all three time series (SO phase, fast and slow sigma power), we extracted the previously identified 2 s windows surrounding each SO trough and concatenated them into segments of 20 SOs, corresponding to 40 s. In case the number of detected SOs was not divisible by 20, the incomplete final segment was padded with SOs randomly resampled from (the same) final segment. The segmentation step was performed to ensure that permutation-based z-scoring (see below) was performed on data segments of identical length, thereby avoiding confounds due to differences in number of detected SOs.

We determined phase-amplitude coupling for each segment of 20 concatenated SOs using an adaptation of the mean vector length method ^61^ that adjusts for possible bias stemming from non-sinusoidal shapes of the modulating frequency ^62^. Specifically, complex-valued debiased phase-amplitude coupling (dPAC) was calculated as:

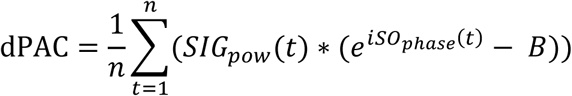
where *i* is the imaginary operator, *t* is time, *SIG_pow_ (t)* and *SO_phase_ (t)* are the sigma power and SO phase time series, and *B* is the mean phase bias:

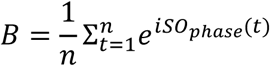

Raw coupling strength (i.e., the degree to which sigma power is non-uniformly distributed across the SO cycle) was defined for each 40 s segment as the magnitude (i.e., length) of the mean complex vector. Under the null-hypothesis of no systematic SO-spindle coupling, vectors of different length should be randomly distributed over SO phases, resulting in a mean vector length close to zero. However, if longer vectors preferentially occur in a particular phase range, the mean vector length will deviate substantially from zero, signaling SO-spindle coupling. Importantly, absolute coupling strength depends on absolute sigma power. Thus, differences in sigma power between electrodes, spindle classes, sleep stages, and individuals, which we have described in detail elsewhere ^22^, confound the interpretation of this measure. Therefore, for every segment and channel, we constructed a surrogate distribution of coupling strengths by repeatedly (n = 1,000) shuffling the SO phase time series with respect to the sigma power time series, and recalculating the mean vector length for each iteration. We then z-scored the observed coupling strength with respect to this null distribution of coupling strength values, and averaged z-scores across segments. Thus, the z-scored measure (dPAC_Z_) indicates how far, in terms of standard deviations, the observed coupling estimate is removed from the average coupling estimate under the null hypothesis of no coupling. Coupling phase (i.e., the SO phase of maximal sigma power) was defined as the phase angle of the mean complex vector, averaged across segments. Control analyses indicated that coupling strength and coupling phase estimates did not depend on the width of the time window surrounding the SO trough (2 vs. 4 s).

### Experimental Design and Statistical Analysis

We examined 58-channel EEG data from 24 healthy volunteers across two consecutive full nights of sleep. While we analyzed both nights, we mainly report analyses from Night 1, except for cross-night comparisons, and analyses involving memory performance, for which (learning) Night 2 was used.

To assess whether coupling strength deviates from chance levels, we compared dPAC_Z_ values to zero with one-sample t-tests. For within-subject comparisons of linear outcome variables (e.g., N3 vs. N2 coupling strength), we used paired-sample t tests. The circular Rayleigh test was used to determine whether circular coupling phase distributions deviate from uniformity. For within-subject comparisons of circular distributions (e.g., N3 vs. N2 coupling phase), we calculated phase differences (wrapped to the −180° to +180° interval) and used a one-sample t-test to compare differences to zero. Associations of coupling phase across nights were assessed using circular-circular correlations, while associations of coupling phase and behavior were tested using circular-linear correlations. Significance of time-frequency power modulations was assessed by performing a one-sample t-test vs. zero at each time-frequency bin. All statistical tests are two-sided, unless stated otherwise. Correction for multiple tests was performed with the False Discovery Rate procedure ^63^.

## Results

We examined oscillatory NREM dynamics in 58-channel EEG data from 24 healthy volunteers across two consecutive full nights of sleep. Night 1 constituted a baseline night, while on Night 2 participants received pre-sleep training on the Motor Sequence Task (MST) and were tested again the next morning. Sleep architecture was in line with typical values for healthy participants and did not differ between nights (Table 1). In what follows, we mainly report findings from Night 1, unless otherwise stated.

**Table 1.** Sleep architecture. Sleep parameters (mean ± SD) in both nights and statistical results from paired t-tests. WASO: wake after sleep onset.

|  | Night 1 | Night 2 | t(23) | P |
| --- | --- | --- | --- | --- |
| N1 (%) | 9.9 ± 5.0 | 10.0 ± 4.4 | -0.1 | 0.95 |
| N2 (%) | 52.0 ± 9.1 | 53.1 ± 7.9 | -0.8 | 0.45 |
| N3 (%) | 19.2 ± 7.8 | 18.5 ± 6.7 | 0.5 | 0.60 |
| REM (%) | 18.9 ± 5.6 | 18.4 ± 6.0 | 0.5 | 0.62 |
| N1 (min) | 48.8 ± 22.7 | 50.1 ± 23.3 | -0.3 | 0.76 |
| N2 (min) | 260.8 ± 57.1 | 265.8 ± 58.0 | -0.4 | 0.67 |
| N3 (min) | 96.1 ± 40.0 | 90.5 ± 31.7 | 0.9 | 0.40 |
| REM (min) | 95.8 ± 33.5 | 90.7 ± 30.1 | 0.8 | 0.41 |
| total sleep (min) | 501.5 ± 61.4 | 497.0 ± 56.5 | 0.3 | 0.73 |
| WASO (min) | 37.1 ± 37.3 | 31.5 ± 16.8 | 0.6 | 0.53 |
| sleep efficiency (%) | 90.3 ± 7.8 | 91.1 ± 4.0 | -0.5 | 0.65 |

### Local dynamics of NREM sleep oscillations

After applying a surface Laplacian spatial filter, we detected SOs on each electrode in both N3 and N2 NREM sleep. Fig. 1A shows a single subject's raw and SO-filtered Laplacian traces for five channels – AFz, FCz, CPz, POz, C6 – at the same time interval, with detected SOs indicated. Sizable channel differences in the SO-filtered traces and their detected SOs are visible, consistent with the notion that most SOs are local phenomena. Similarly, slow and fast sigma activity showed notable differences between channels, indicating that spindle dynamics, like SOs, show regional variability. Substantial within-channel differences in the time courses of slow and fast sigma are also apparent, suggesting that the two sigma bands reflect different neuronal dynamics. This clear evidence of regional variation within spindle and SO frequency bands motivated us to examine channel-specific SO-spindle coupling.

**Figure 1.**
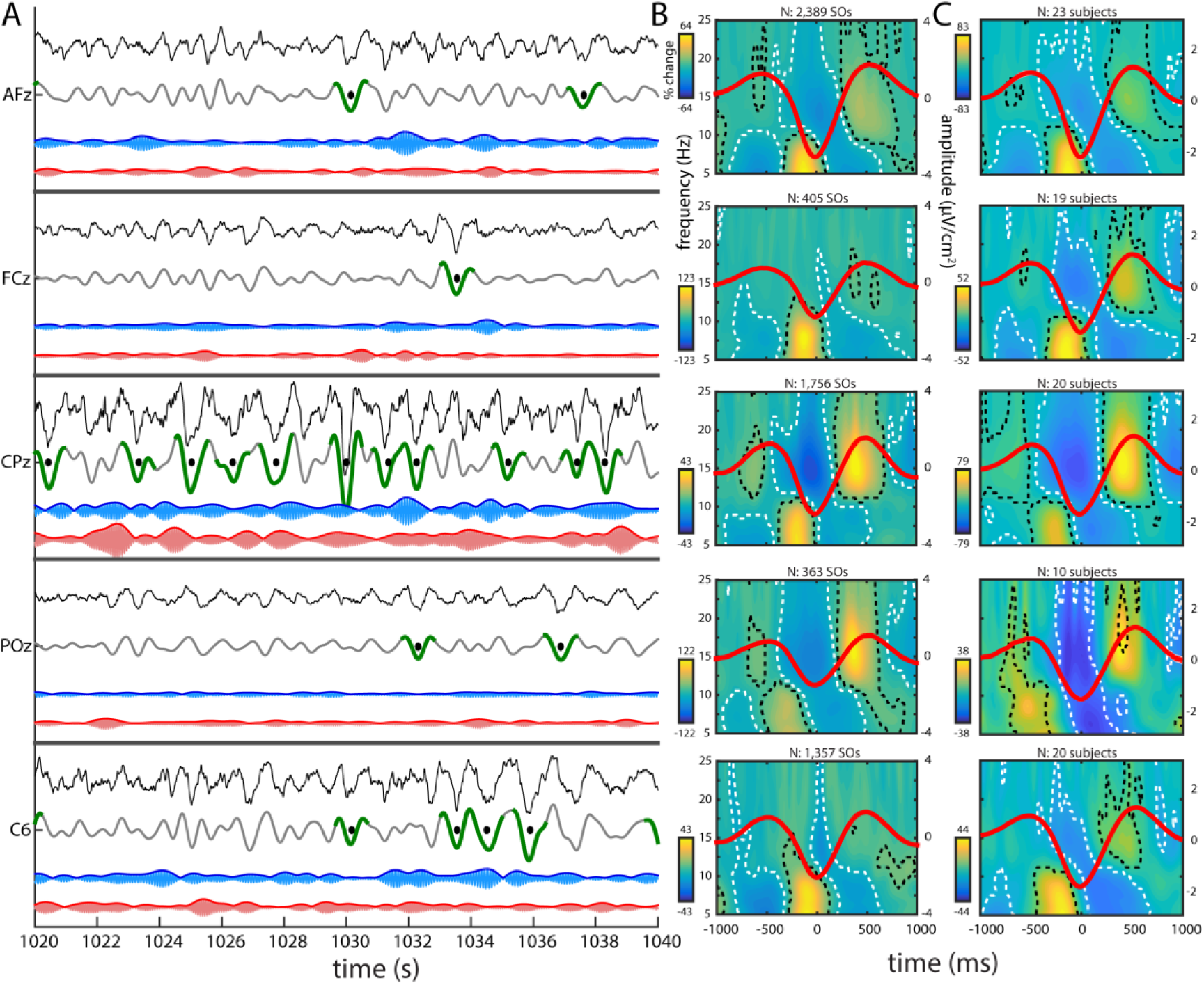
Regional variations in SO, slow spindle, and fast spindle activity. (A) Twenty second excerpt of single subject's N3 sleep for five channels - AFz, FCz, CPz, POz, C6 - showing, from top to bottom, the raw Laplacian EEG (black), SO- (gray), slow sigma- (blue), and fast sigma-filtered (red) traces. Detected SOs are shown in green (trough ± 500 ms), with the troughs marked with black dots. Sigma traces were multiplied by a factor 6 for visualization purposes. (B) Same subject's time-frequency spectrograms time-locked to SO troughs (average SO waveforms superimposed in red; total number of detected SOs above spectrogram). Activity (percentage power change relative to baseline; see Methods) in the slow sigma/theta and fast sigma ranges is modulated by SOs on each channel, as signified by clusters of significant power increases and decreases (indicated by black and white dashed outlines, respectively). Significance was assessed by performing a one-sample t-test vs. zero at each time-frequency bin, followed by False Discovery Rate correction for multiple comparisons (P_adj_<0.05). (C) Group-level time-frequency spectrograms time-locked to SO troughs. Subjects included only when ≥ 20 SOs. Other details as in (B).

To illustrate local SO-spindle coupling for this sample subject using conventional techniques, we time-locked each sample channel's raw Laplacian time series to that channel's identified SO troughs, allowing us to evaluate time-frequency power in the 5–25 Hz range as a function of the SO waveform. This revealed significant modulations of faster activity on each sample channel (Fig. 1B), similar to previous reports detecting global ^30,32^ or local SOs ^31^. Specifically, we found robust increases in ~15 Hz fast sigma power, extending to higher frequencies, centered on the SO peaks preceding and following the SO trough, while fast sigma was suppressed in the SO trough. In contrast, activity in the slow sigma and theta range (~5–10 Hz) was markedly enhanced prior to the SO trough, but suppressed in the SO peaks preceding and following the SO trough. Similar observations were made for the majority of individual channels and subjects, as well as across subjects (Fig. 1C). Thus, despite different temporal dynamics of SO and sigma activity on different channels, the relation between local SO phase and local sigma power appears to remain intact.

While the time-frequency approach demonstrates local coupling, this method suffers from limited temporal and spectral precision. It cannot specify the precise SO phase at which activity in the fast and slow sigma ranges peaks, necessitating alternative approaches, as described in subsequent sections.

### SO characteristics

Overall, across subjects and sleep stages on Night 1, we detected 753,641 SOs. Although subjects spent far less time in N3 than N2 (Table 1), almost four times as many SOs were detected in N3 as in N2 (20,089 ± 19,232 vs. 5,729 ± 4,943, t(23)=3.9, P<0.001). Corresponding channel-averaged SO densities (number per minute) showed a similarly significant six-fold sleep stage difference (N3: 2.4 ± 1.5; N2: 0.4 ± 0.3; t(23)=7.0, P<10^−6^). Topographical examinations (Fig. 2A) confirmed this stage difference while also revealing known regional differences in the prevalence of SOs, with markedly higher SO densities over anterior and central electrodes than in temporal and posterior areas ^64^. SO densities for the five sample channels are shown in Table 2 for both nights. Trough-to-peak SO amplitudes were greatest at frontal electrodes, but relatively uniform over the rest of the scalp (Fig. 2B). Averaged across electrodes, SOs were of slightly but significantly larger amplitude in N3 (2.84 ± 0.26 μV/cm^2^) relative to N2 (2.76 ± 0.22 μV/cm^2^; t(23)=2.2, P=0.04). SO characteristics were very similar for Night 2.

**Figure 2.**
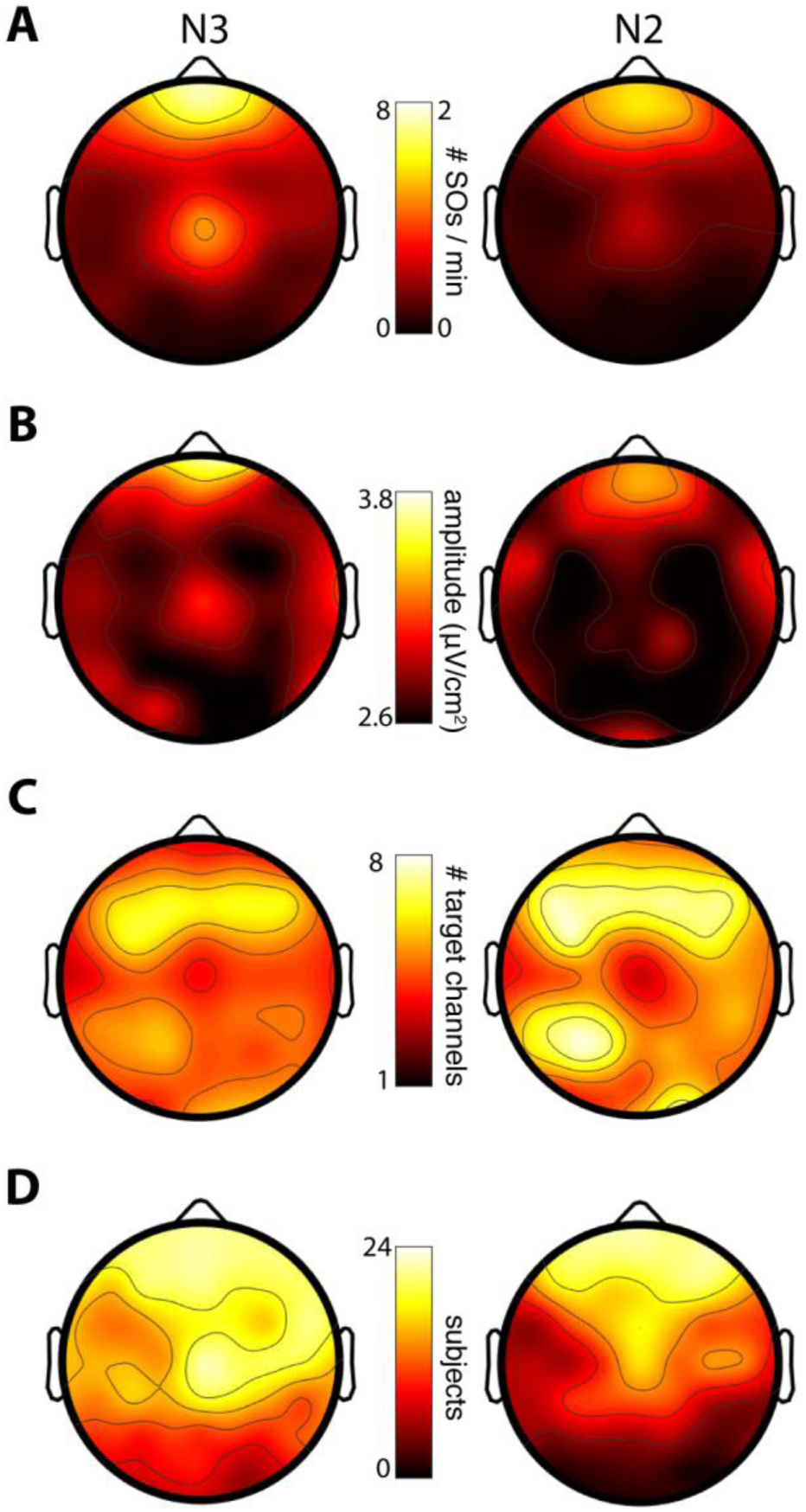
Topographical distributions of slow oscillations in N3 and N2. (A) SO densities (number/minute). Note the different scales for N3 and N2 SO density. (B) SO amplitudes (trough-to-peak). (C) Average number of target channels with an SO trough co-occurring within 100 ms surrounding the SO trough of a source channel. (D) Number of subjects included.

**Table 2.** SO densities for sample channels. SO densities (mean ± SD) in both nights and statistical results from paired t-tests.

| channel | stage | Night 1 | Night 2 | t(23) | P |
| --- | --- | --- | --- | --- | --- |
| Afz | N3 | 7.1 ± 6.0 | 7.6 ± 6.8 | -0.5 | 0.64 |
|  | N2 | 1.5 ± 1.1 | 1.5 ± 1.1 | -0.2 | 0.84 |
| FCz | N3 | 2.8 ± 2.6 | 2.4 ± 2.2 | 0.7 | 0.51 |
|  | N2 | 0.5 ± 0.5 | 0.5 ± 0.5 | 0.2 | 0.84 |
| CPz | N3 | 4.1 ± 4.1 | 3.1 ± 3.0 | 1.3 | 0.22 |
|  | N2 | 0.4 ± 0.4 | 0.4 ± 0.5 | -0.2 | 0.88 |
| Poz | N3 | 0.6 ± 1.0 | 0.7 ± 1.2 | -0.4 | 0.70 |
|  | N2 | 0.1 ± 0.1 | 0.1 ± 0.2 | -0.9 | 0.39 |
| C6 | N3 | 2.0 ± 2.1 | 1.5 ± 2.2 | 0.9 | 0.38 |
|  | N2 | 0.3 ± 0.5 | 0.2 ± 0.4 | 1.1 | 0.26 |

### SO co-occurrences

To formally assess the degree to which SOs are local, we determined the number of "target" channels that showed SOs co-occurring with each SO detected on a "source" channel. We counted a co-occurrence when the SO trough of a target channel occurred within a fixed window surrounding the source SO trough. Using a liberal window of ± 400 ms on either side ^20^, we found that SOs involved fewer than 10 channels on average (mean number of target channels across source channels, averaged across SOs and subjects; N3: 8.6 ± 1.3; N2: 9.4 ± 1.5), indicating that most detected SOs engage only a limited number of electrodes. Despite their large difference in SO densities, N3 and N2 SOs showed very similar numbers of co-occurring channels, although channel involvement was significantly higher in N2 than N3 (t(57)=5.6, P<10^−6^).

Next, we narrowed the window of co-occurrence to ± 100 ms in order to count only co-occurrences where SOs showed minimal phase shifts across channels. This led to a 40–45% reduction in the average number of channels participating in each SO, leaving 4.9 ± 0.9 and 5.7 ± 1.2 involved channels in N3 and N2, respectively (paired t-tests for wide vs. narrow window: both t(57)>22.2, P<10^−29^; stage difference: t(57)=9.0, P<10^−11^). In other words, only roughly half of target-channel SOs occurring in conjunction with a source SO were approximately in phase with the source SO. Further topographical examinations (Fig. 2C) using this restricted window width indicated greatest co-occurrence for SOs detected in anterior source channels, with this hotspot shifted slightly posteriorly relative to the sites with highest SO density and amplitude.

The preceding analyses are based on mean channel involvement averaged across all SOs detected in each source channel. We next examined the number of involved channels across individual SOs (using the restricted 100 ms window). Normalized histograms for two sample channels, averaged across subjects, illustrate the skewedness of these distributions, with the majority of SOs affecting only a minority of channels, with the modal number of channels involved being only 1 and 2 (Fig. 3). Across source channels, the modal number of target channels ranged between 1 and 11 (N3: 2.7 ± 1.7; N2: 2.6 ± 1.9; t(57)=0.4, P=0.69), with greatest channel involvement again seen for frontal source channels. Cumulative distributions indicated that, on average across source channels, 50% of SOs involved no more than 10% of channels (mean number of target channels for N3: 4.3 ± 0.9; N2: 5.0 ± 1.5; t(57)=4.7, P<10^−4^). Moreover, 75% of all SOs were detected in no more than 15% of channels (N3: 7.3 ± 1.1; N2: 8.7 ± 2.2; t(57)=6.2, P<10^−7^), and less than 1% of SOs involved more than a third of recording sites (N3: 15.9 ± 1.5; N2: 18.6 ± 2.6; t(57)=8.6, P<10^−11^). In sum, this pattern of results, which was reproducible across nights, indicates that SOs are overwhelmingly local in nature (especially in N3).

**Figure 3.**
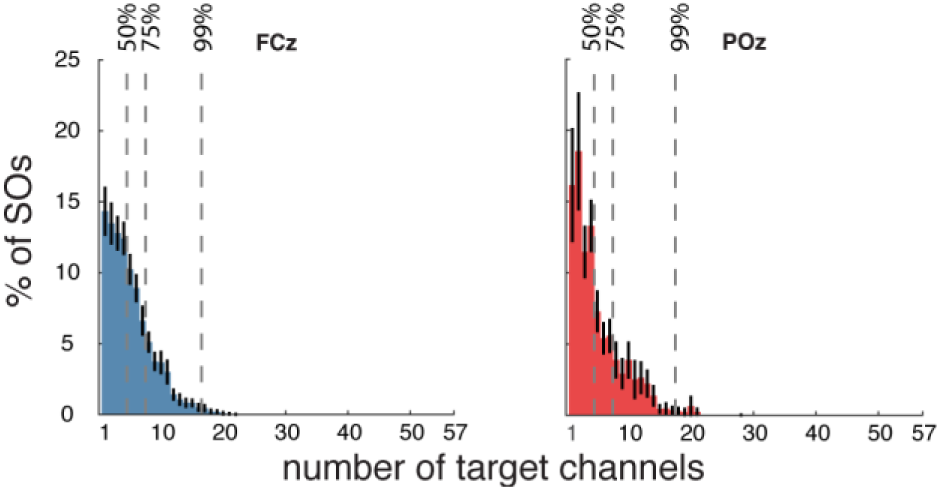
Distribution of SO involvement (percentage of individual source SOs detected on a given number of target channels). Mean ± SEM (black error bars) across subjects for N3 SOs detected on anterior (FCz) and posterior (POz) source channels. Vertical dashed lines indicate channel count where cumulative distribution contains 50, 75, and 99% of SOs. Distributions were qualitatively similar for all channels and sleep stages.

There were substantial individual differences in the number and density of detected SOs, with some subjects expressing few or no SOs meeting our detection criteria on one or more channels. To attain reliable estimates of SO-spindle coupling in subsequent analyses, we included, for each electrode, only those subjects showing a minimum of 20 detected SOs (evaluated separately for N3 and N2). Using this criterion, we included an average of 16.0 ± 5.1 subjects per electrode (range: 5–23) in N3, and 11.1 ± 7.1 (range: 1–23) in N2. Conversely, we included 38.8 ± 14.1 electrodes per subject (range: 4–57) in N3, and 26.8 ± 12.6 (range: 11–56) in N2. Fig. 2D displays the number of subjects included at each electrode, showing reduced inclusion rates over posterior areas, especially during N2.

### SO-spindle coupling

To determine cross-frequency coupling between locally detected SOs and local spindle activity we employed the debiased phase-amplitude coupling (dPAC) approach ^62^. The dPAC method provides information on both coupling strength (i.e., the degree to which sigma activity is non-uniformly distributed across the SO cycle), and coupling phase (i.e., the SO phase of maximal sigma activity). Using this method, we determined SO-spindle coupling at each electrode, separately for N3 and N2 sleep, and separately for fast and slow sigma activity. Fast and slow sigma ranges were set for each individual based on their own power spectrum (fast sigma: 13.5 ± 0.6 Hz; slow sigma: 10.9 ± 0.7 Hz), thereby targeting spindle activity in a subject-specific manner ^22^. We further normalized coupling strengths using a permutation-based reshuffling approach, resulting in z-scored coupling strengths (dPAC_Z_). Critically, this normalized measure of coupling strength is independent of absolute sigma power and is therefore not influenced by potential differences in sigma power between electrodes, sleep stages, or individuals.

### Local SOs coordinate local spindle activity across the scalp

We first sought to determine the presence of SO-spindle coupling (i.e., above-zero coupling strengths), irrespective of coupling phase. We did this separately for each combination of fast/slow spindles and N3/N2 sleep (henceforth: conditions). In a first step, we averaged coupling strengths across subjects for each electrode. Average channel coupling strengths were significantly greater than zero in each condition (all t(57)>39.2, P<10^−42^). Although absolute differences were small, coupling was significantly stronger in N3 than N2 for fast sigma, while the reverse was the case for slow sigma (Table 3). Coupling strengths were also significantly greater for fast than slow sigma in N3, but not different in N2.

**Table 3.** SO-spindle coupling strengths. Mean (± SD) coupling strengths (dPAC_Z_) in each condition and statistical results from paired t-tests.

|  | N3 | N2 | t(57) | P |
| --- | --- | --- | --- | --- |
| fast | 29.2 ± 5.1 | 25.7 ± 5.0 | 4.8 | <10<sup>-4</sup> |
| slow | 23.5 ± 2.3 | 26.2 ± 3.9 | 4.7 | <10<sup>-4</sup> |
| t(57) | 7.7 | 0.6 |  |  |
| P | <10<sup>-9</sup> | 0.55 |  |  |

To assess the spatial extent of this SO-spindle coupling, we next analyzed coupling strengths separately for each electrode. Correcting for multiple comparisons with the False Discovery Rate (FDR) procedure ^63^, all 58 electrodes showed significant (P_adj_<0.05) SO-spindle coupling during N3 for both fast and slow sigma, while 52 of 58 (fast) and 51 of 58 (slow) electrodes reached significance for N2. The few electrodes not reaching significance were exclusively positioned over posterior regions with low numbers of both SOs and included subjects.

We further assessed the proportion of included (≥ 20 SOs) subject-electrodes that exhibited evidence of SO-spindle coupling, operationalized as dPAC_Z_ values > 1.65 (corresponding to one-sided P<0.05, uncorrected). Across N3 (n = 930) and N2 (n = 642), all 3,144 subject-electrode-condition combinations showed significant coupling except for a single subject-electrode that failed to reach significance for fast spindle coupling during N3 (99.97%). Highly similar results were found for Night 2. These analyses confirm the robustness of SO-spindle coupling at all scalp sites, for both fast and slow spindles, and in both N3 and N2 sleep.

### Local coupling phase depends on spindle class and sleep stage

Given this widespread coupling of local spindle activity to local SOs, we turned to topographical analyses of coupling phase (i.e., the SO phase at which sigma activity is greatest). For fast spindles, activity was preferentially expressed on the rising slope of the SO in both N3 (group averages across electrodes: 50 ± 29°; Fig. 4A) and N2 (80 ± 33°; Fig. 4B; see next paragraph for direct stage comparisons). Phase distributions across channels were highly non-uniform (Rayleigh test: both P<10^−22^; see dashed insets), reflecting the consistency of coupling phases across the scalp. To assess the consistency of coupling phases across subjects, we examined, for each electrode, whether coupling phases were non-uniformly distributed. Right panels of Fig. 4AB shows these distributions for the five sample channels, and indicate tight group-level phase clustering before the SO peak on most channels (although clear between-subject differences are evident even for the most consistent electrodes; see below). Across the scalp, 46/58 (N3; uncorrected: 49/58) and 26/58 (N2; uncorrected: 32/58) electrodes showed significantly non-uniform distributions of coupling phase, indicating highly consistent group effects over much of the scalp (significant electrodes indicated on topographies as green circles). The failure of most posterior channels to reach significance is most like due to the small number of subjects with sufficient numbers of SOs to be included in the analysis, particularly in N2.

**Figure 4.**
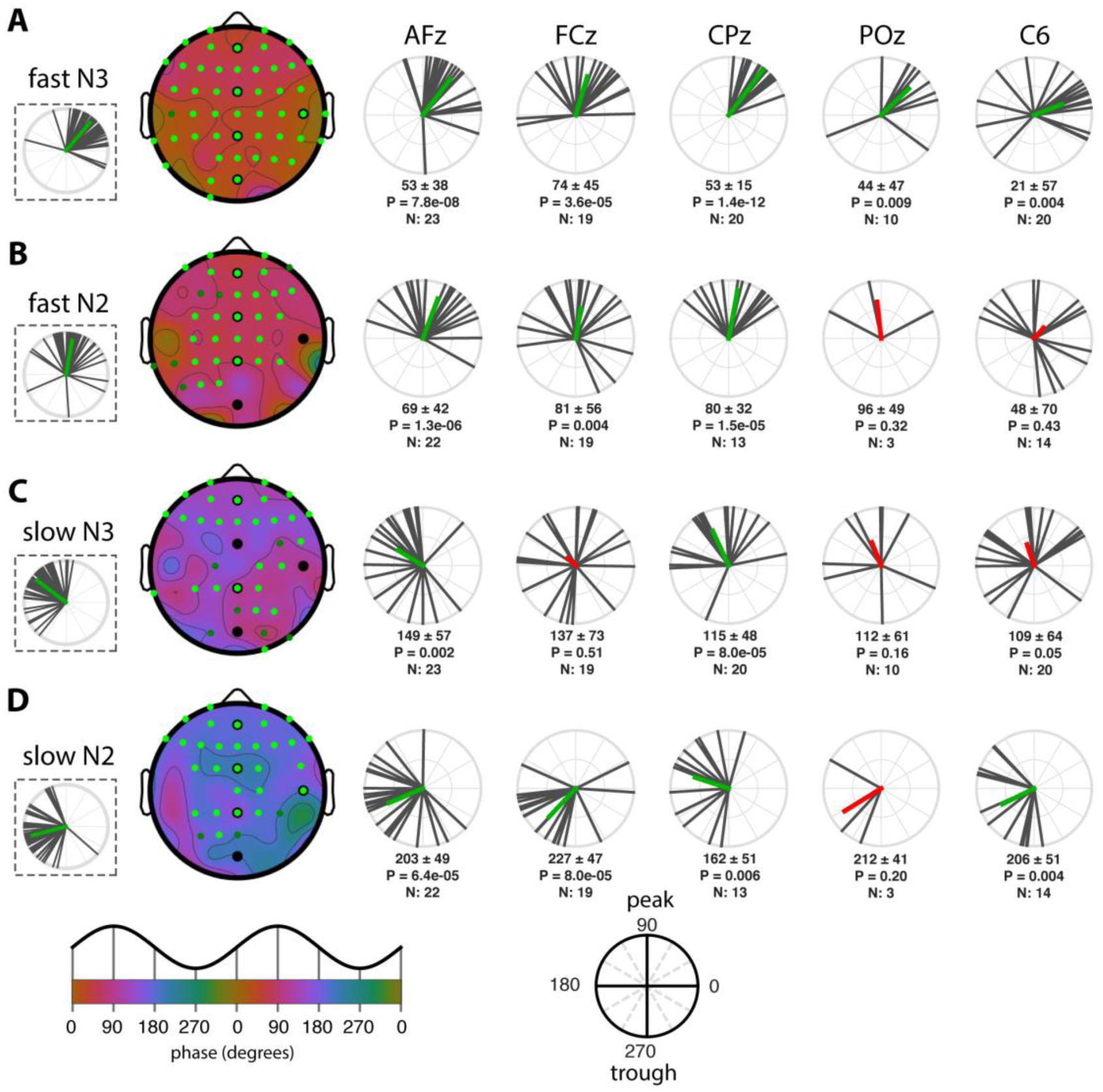
SO-spindle coupling phases across the scalp and individuals. (A) fast N3 spindles. (B) fast N2 spindles. (C) slow N3 spindles. (D) slow N2 spindles. **Left:** topographies of mean preferred coupling phase across individuals, with dashed insets on the left showing circular distributions across channels (black vectors: individual channels, averaged across individuals; green vectors: channel averages). Green colored circles on topographical maps indicate electrodes showing significantly non-uniform phase distributions across subjects after correcting for multiple comparisons (bright green) or uncorrected (dark green). Black circles indicate selected electrodes plotted on the right. **Right:** circular plots display phase distributions across individuals (black). Subject-average vectors (colored) indicate both average phase and cross-subject consistency, expressed as its length. Numbers below each circular plot indicate mean phase (± SD), P value (uncorrected) of Rayleigh tests for non-uniformity, and number of included subjects. Mean vector is colored green when the uncorrected P is <0.05, and red otherwise. Bottom insets indicate mapping from phases to topographical color map and circular plots.

Next, we asked whether fast spindle activity peaked in different SO phases in N3 and N2 by calculating within-subject N3–N2 phase differences at each electrode and comparing these to zero. Subjects were included only if they had ≥ 20 SOs in both N3 and N2. Across the scalp, averaged within-subject N3–N2 coupling-phase differences were significantly different from zero (−15 ± 37°; t(57)=2.5, P=0.02), with fast spindles occurring slightly earlier in N3 than N2. (Note that this estimate of within-subject phase difference differs from the 30° difference based on group averages seen in the previous paragraph.) This small (15°) difference in preferred coupling phase, together with the fact that no individual electrode showed a significant N3–N2 difference after multiple comparison correction (uncorrected: 8/58), leads us to conclude that the organization of fast spindle activity by SOs does not differ importantly between these sleep stages.

Slow spindles were organized by SOs quite differently than fast spindles, showing maximal activity in the transition from the SO peak to the trough (group averages across electrodes: N3: 143 ± 30°, Fig 4C; N2: 195 ± 34°, Fig. 4D). As for fast spindles, phase distributions were highly non-uniform (Rayleigh: N3 and N2, both P<10^−21^), indicating that slow spindles are preferentially expressed in similar SO phases across the scalp in both N3 and N2. Examining cross-subject consistency, we found that 27/58 (N3; uncorrected: 33/58) and 29/58 (N2; uncorrected: 32/58) electrodes showed significantly non-uniform phase distributions across subjects, with anterior regions expressing this most consistently across sleep stages. Although indicative of consistent group-level clustering over much of the scalp, these findings and the circular distributions of Fig. 4CD indicate that between-subject variability of coupling phases is generally greater for slow than fast spindle activity.

Examining N3–N2 stage differences for slow-spindle coupling, we found that within-subject phase differences were significantly different from zero (group average across electrodes: −48 ± 37°; t(57)=9.0, P<10^−11^), with the phase of maximal slow-spindle coupling about 45° earlier in N3. Testing of individual electrodes followed by multiple comparison correction revealed that of the 58 electrodes, 12 electrodes (7 anterior, 5 central; uncorrected: 14) had significantly different coupling phases in N3 compared to N2 (e.g., circular distributions of AFz in Fig. 4CD).

Finally, we directly compared the preferred coupling phases of fast and slow spindles. Within-subject fast-slow phase differences were significant in both N3 (group average across electrodes: fast-slow = −80 ± 42°; t(57)=7.0, P<10^−8^; Fig. 5A), and N2 (−114 ± 45°, t(57)=−6.2, P<10^−7^; Fig. 5B), indicating that maximal fast spindle activity occurs about a quarter of a cycle earlier than for slow spindles. The larger phase difference in N2 is consistent with the aforementioned observation of slow spindles being phase-delayed in N2 relative to N3. On an electrode-by-electrode basis, fast-slow differences were confirmed statistically for 17/58 electrodes in N3, and 2/58 in N2 using the most stringent correction for multiple comparisons (Fig. 5, green circles). However, using an uncorrected threshold of P<0.05 increased the channel count to 22 of 58 in N3 and 18 of 58 in N2 (Fig. 5, green + white circles). In sum, these findings indicate that fast spindle activity is expressed distinctly earlier in the SO cycle than is slow spindle activity, and that this phenomenon can be observed across widespread cortical regions, particularly during N3.

**Figure 5.**
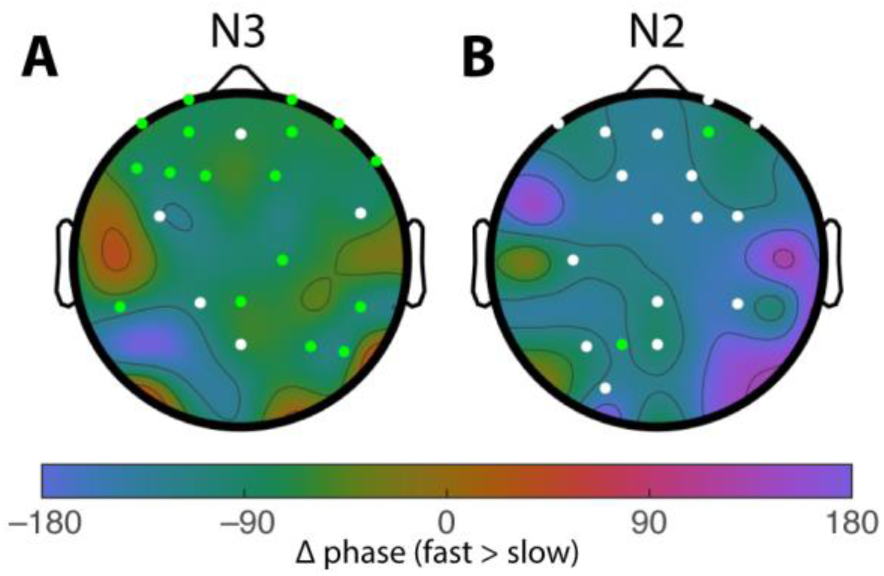
Differences in coupling phase between fast and slow spindle activity. Fast spindles occur ~90° earlier than slow spindles in both N3 (A) and N2 (B). Colored circles indicate electrodes showing significant phase differences across subjects after FDR correction (green) or uncorrected (white).

Combined, these findings, which were similar for Night 2, indicate that fast and slow spindle activity are tied to different phases of the local SO cycle, but also that N3 and N2 SOs coordinate spindle activity in subtly different ways.

### Regional differences in coupling phase

While the preceding analyses demonstrated relatively consistent coupling phases across the scalp for all combinations of spindle class and sleep stage, the circular distributions of Fig. 4A (insets) also indicate substantial between-channel variability. To determine if there was any systematic pattern to this, we assigned channels to one of three regions (anterior, central, posterior; Fig. 6, inset) and averaged coupling phases across channels in each region for each subject. The resulting distributions were all significantly non-uniform (all P_adj_<0.05), indicating consistent group-level clustering of coupling phase within each region for each condition. Also visible in these plots are the previously identified differences in coupling phase across conditions.

Considering regional differences, N3 spindles appeared to peak around 25° later in the SO cycle in anterior compared to central and posterior regions (Fig. 6AC), while no differences were apparent during N2 (Fig. 6BD). To evaluate these observations statistically, we determined interregional phase differences separately for each subject to account for individual differences not captured by the group-level approach of Fig. 6, and compared the resulting values to zero (one-sample t tests). These analyses confirmed that anterior N3 spindles occurred significantly later in the SO cycle compared to central (for fast and slow spindles) and posterior (for fast spindles) areas, although these analyses did not survive multiple comparison correction (detailed statistics in Table 4). However, analyses from Night 2 confirmed the anterior vs. central difference for both fast and slow N3 spindles, thus suggesting subtle regional differences regarding the SO phase of maximal spindle expression. Although it might be expected that the degree of cortical depolarization/hyperpolarization, as approximated by the trough-to-peak SO amplitude, explains this regional variability, N3 SO amplitudes did not differ reliably between anterior and central regions (Night 1: P_adj_=0.29; Night 2: P_adj_=0.32).

**Figure 6.**
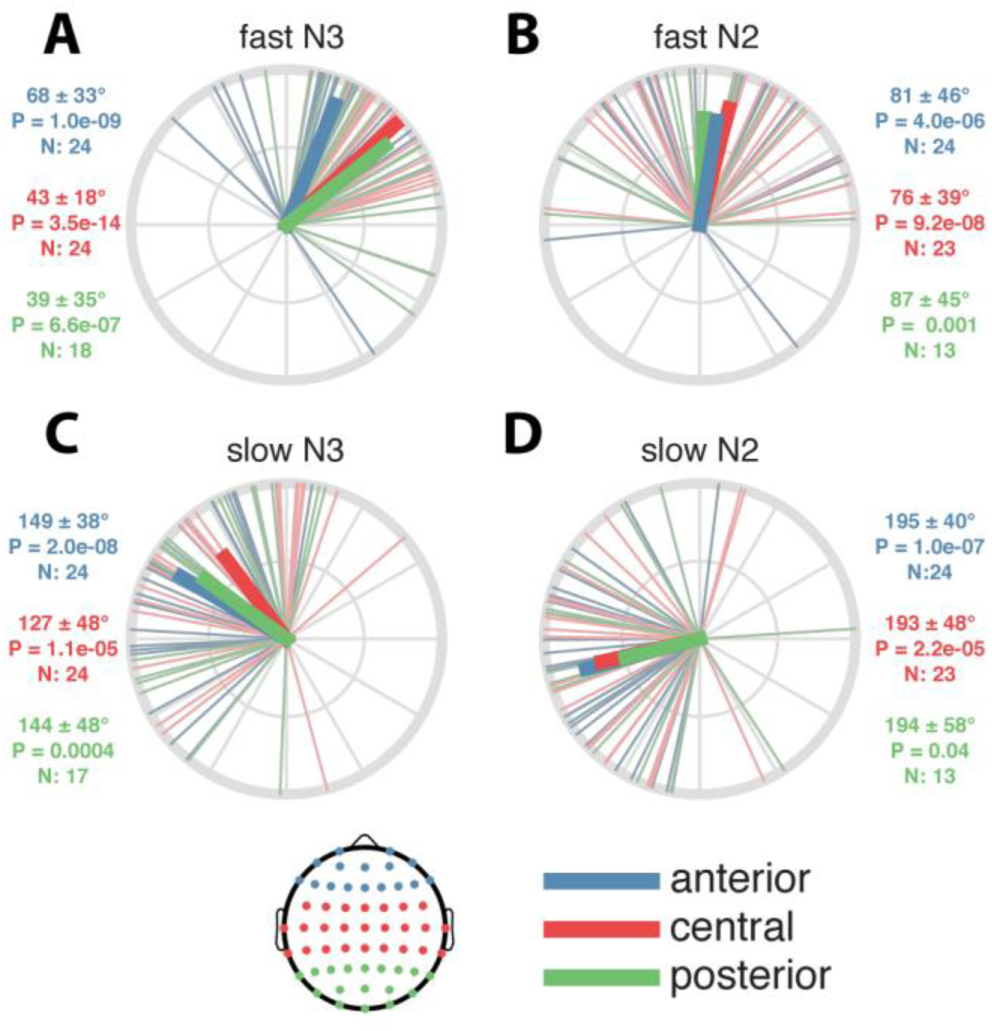
Regional differences in coupling phase. Numbers next to each circular plot indicate mean phase (± SD), P value (uncorrected) of Rayleigh test for non-uniformity, and number of included subjects. Inset: topography showing assignment of channels to regions.

**Table 4.**
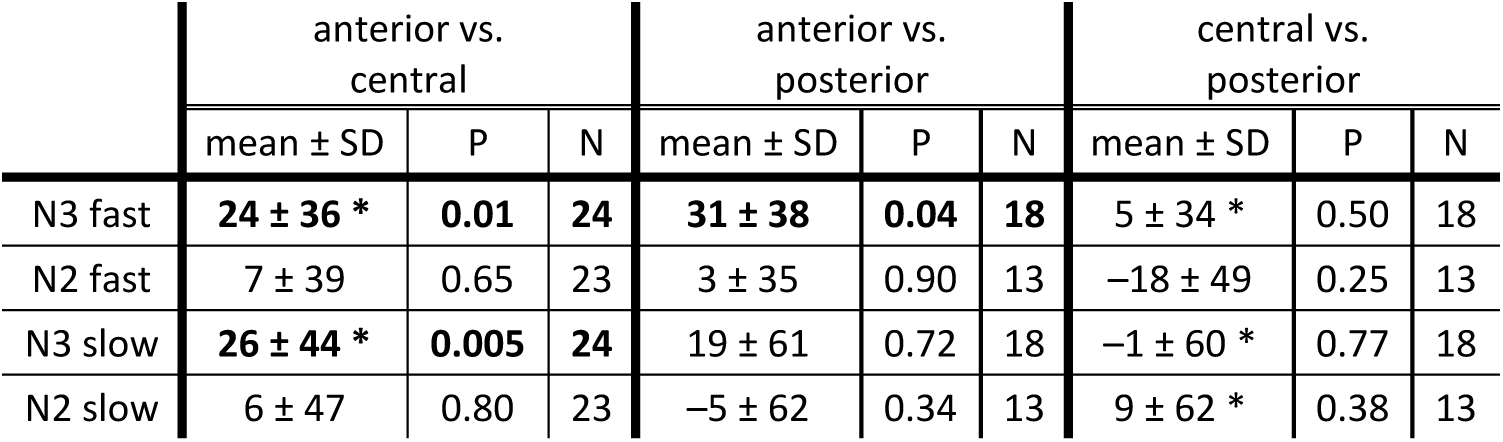
Regional differences in coupling phase. Indicated are mean phase difference (± SD), P value (uncorrected) of one-sample t test vs. zero, and number of included subjects. Note that for these analyses, subjects are included only when they have ≥ 20 SOs on at least one electrode in each of the compared regions. Bold entries: significant phase difference on Night 1 (shown in Fig. 6). Asterisks: significant phase difference on Night 2.

|  | anterior vs. central |  |  | anterior vs. posterior |  |  | central vs. posterior |  |  |
| --- | --- | --- | --- | --- | --- | --- | --- | --- | --- |
| | mean ± SD | P | N | mean ± SD | P | N | mean ± SD | P | N |
| N3 fast | **24 ± 36 \*** | **0.01** | **24** | **31 ± 38** | **0.04** | **18** | 5 ± 34 * | 0.50 | 18 |
| N2 fast | 7 ± 39 | 0.65 | 23 | 3 ± 35 | 0.90 | 13 | -18 ± 49 | 0.25 | 13 |
| N3 slow | **26 ± 44 \*** | **0.005** | **24** | 19 ± 61 | 0.72 | 18 | -1 ± 60 * | 0.77 | 18 |
| N2 slow | 6 ± 47 | 0.80 | 23 | -5 ± 62 | 0.34 | 13 | 9 ± 62 * | 0.38 | 13 |

### Individual differences in coupling phase are stable across nights

Beside regional differences in coupling phase, the circular plots of Fig. 4 (right) and Fig. 6 also indicate substantial individual differences in the exact phase of coupling. To further examine this phenomenon, we analyzed the distribution of coupling phases across channels within subjects. As illustrated in the top row of Fig. 7 for two sample subjects, both individuals showed highly non-uniform distributions across channels with fast spindles (Fig. 7A) most prominent preceding, and slow spindles (Fig. 7B) following the SO peak. However, coupling phases clearly differed between these individuals, for both fast and slow spindles. Interestingly, plotting the same subjects' phase distributions for their second night indicated that individual differences in coupling phase are consistent across nights (Fig. 7AB, bottom), raising the possibility that this variability constitutes a stable trait.

**Figure 7.**
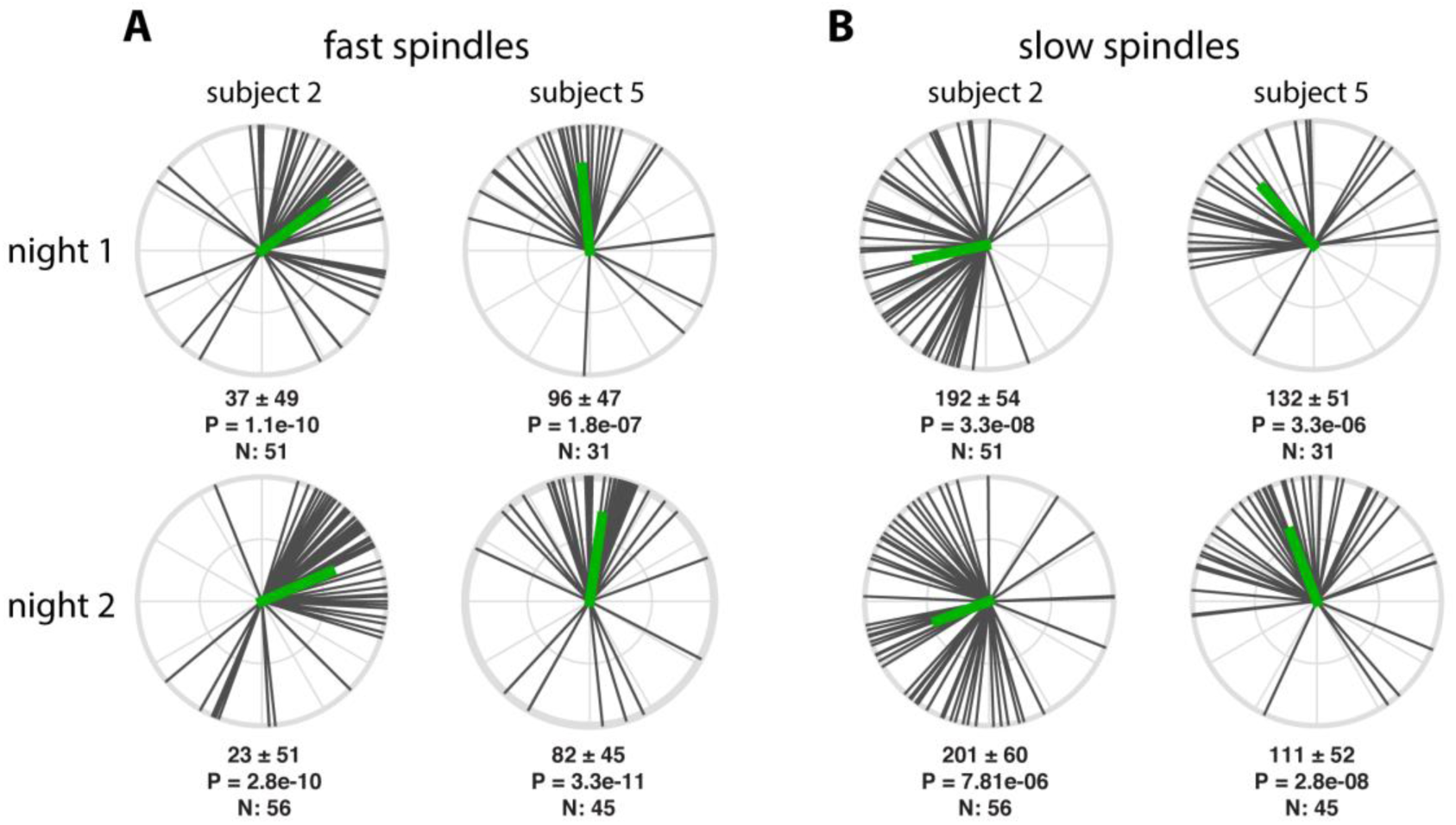
Individual variability of N3 coupling phases for two example subjects. Numbers below each circular plot indicate mean phase across channels (± SD), P value (uncorrected) of circular tests for non-uniformity, and number of included electrodes.

To further examine this notion, we used circular correlation techniques to determine if there is a reliable association between subjects' coupling phases across nights. Given the general consistency of coupling phases across channels, we first averaged coupling phases across all available channels for each subject. Using this approach, we obtained remarkably strong cross-night correlations of coupling phase in each condition (all P_adj_<0.04), indicating that individuals who express spindles in a later SO phase in one night tend to show the same pattern in the next night (Fig. 8). This was the case even for fast N3 spindles, where strong group-level clustering left relatively little between-subject variance (phases Night 1: 52 ± 19°; span: 92°). In contrast, between-subject phase variability was much greater for fast N2 spindles (79 ± 46°; span: 203°), and slow spindles in both N3 (141 ± 35°; span: 153°;) and N2 (194 ± 36° span: 162°), spanning approximately half of the SO cycle. Thus, some subjects express spindles in an SO phase that deviates substantially from the group average, but they do so consistently across nights. We examined whether the observed variability in coupling phase could be explained by differences in trough-to-peak SO amplitude. However, circular-linear correlation analyses did not indicate a relation between these variables in any condition in Night 1 (all P_adj_>0.29), or Night 2 (all P_adj_>0.15). We did not find evidence that coupling dynamics differed between the two nights (one-sample t tests of within-subject phase differences vs. zero: all P>0.18), suggesting that there were no task-induced alterations of SO-spindle coupling dynamics.

**Figure 8.**
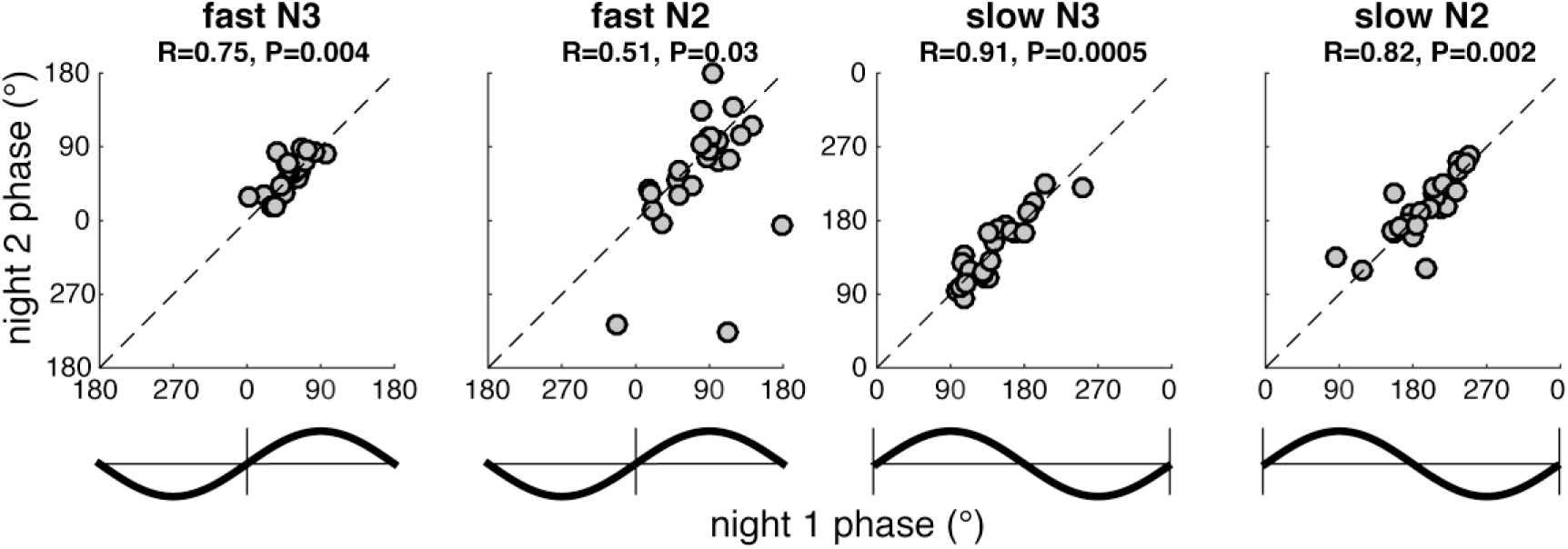
Stable individual differences of global SO-spindle coupling phase. Each scatter plot shows individuals' preferred coupling phases for nights 1 (x-axis) and 2 (y-axis). Each individual's coupling phase estimates are averaged across all available electrodes. Correlation coefficients (R) and (uncorrected) P values from circular correlation analyses are indicated above each plot. Note the different axis scales for fast and slow spindles, also indicated by schematic SO waveforms. Also note that because of the circular nature of phase estimates, data points close to the edge "wrap around" and are also close to the opposite side.

### Cross-night stability of regional SO-spindle coupling

Given that we observed significant regional variability of coupling phases within subjects (Fig. 6; Table 4), we repeated the preceding cross-night correlations separately for anterior, central, and posterior regions (Fig. 9). These analyses indicated that individual differences in coupling phase were significantly correlated across nights in anterior and central regions for three out of four conditions (all P_adj_<0.05), with the remaining correlations showing trends to significance. In contrast, posterior coupling phases showed a significant correlation only for fast N3 spindles, although this association did not survive correction for multiple comparisons. However, we emphasize that an absence of significant cross-night correlations in posterior areas does not imply that spindles are not consistently coordinated by the regional SO cycle, only that there is more night-to-night variability within subjects. This observation may again be related to the smaller number of detected SOs and included subjects for posterior analyses. Also note how observations are clustered into different SO phases for different regions, spindle classes, and sleep stages, consistent with the observations presented in previous sections. As for the global analyses reported in the previous section, individual differences in coupling phase were not related to between-subject variability in SO amplitude within regions, for neither Night 1 (all P_adj_>0.21), nor Night 2 (all P_adj_>0.29).

**Figure 9.**
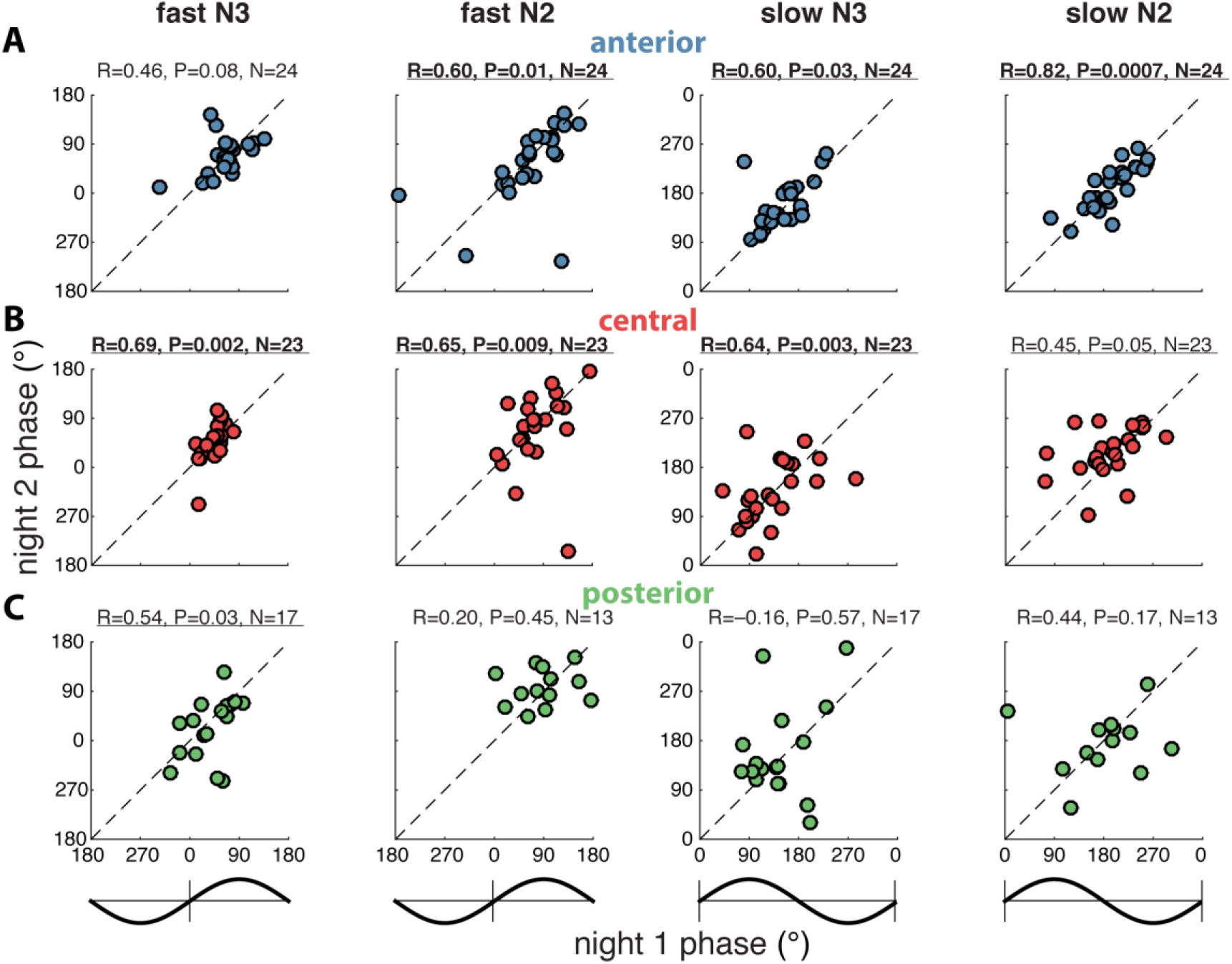
Stable individual differences of regional SO-spindle coupling phase. Figure layout as in Fig. 8, only with coupling phases averaged separately across available anterior (A), central (B), and posterior electrodes (C). Number of included subjects (N) indicated above each plot. Note that for these analyses, subjects are included only when they have ≥ 20 SOs on at least one electrode in the analyzed region in both nights. Significant (P<0.05, uncorrected) correlations are underlined, and correlations surviving multiple comparison correction are indicated in bold.

### SO-spindle coupling and memory

Given previous reports that SO-spindle dynamics may be predictive of overnight memory retention ^16,38,39^, we explored whether individual variability in coupling phase on Night 2 (i.e., the learning night) is associated with procedural memory change. Individuals showed significant improvement in the number of correctly completed motor sequences on the MST (16.3 ± 12.6%; t(23)=6.4, P<10^−5^), consistent with typical overnight gains in healthy subjects ^38,50–52^. However, we did not observe reliable associations between the SO phase of preferential spindle expression and memory improvement, for either fast or slow spindles, in either N3 or N2 sleep, and using either global or regional (as defined in the preceding two sections) estimates of coupling phase (circular-linear correlation analyses: all P>0.11).

## Discussion

The current study addresses the large-scale dynamics of SO-spindle coupling. Our main findings are that 1) local spindle activity is coupled to the local SO phase; 2) spindles are preferentially expressed in distinct phases of the SO cycle as a function of a) spindle class (slow vs. fast); b) sleep stage; and c) cortical region; and 3) individual differences in coupling phase are stable across nights.

We found that the phase of locally detected SOs robustly orchestrates local spindle activity at virtually every electrode and across all combinations of spindle class and sleep stage (Table 3), extending earlier evidence of regionally restricted spindle modulation by the local SO phase ^31^. These findings are relevant to theories that SO-spindle coupling facilitates the recoding of temporary memory traces to more permanent cortical representations ^65,66^. Specifically, the demonstration that local SO-spindle coupling extends across the cortex adds an important spatial component to this temporal coordination, providing the neurophysiological prerequisite for circuit-specific plasticity processes and the consolidation of specific memory traces ^33,47–49^.

Regarding the precise SO phase of maximal spindle activity, we observed both known and novel sources of variability in coupling dynamics. First, our findings confirm the general dissociation of scalp-recorded fast and slow spindle activity, with fast spindles having maximal amplitude preceding, and slow spindles following, the SO peak ^30–33^ (but see ^34,36^ for no apparent slow/fast spindle differences in intracranial studies). This phenomenon was observed across the scalp for both N3 and N2 (Fig. 4), with fast spindles occurring approximately a quarter of a cycle earlier, most robustly in anterior areas (Fig. 5). These findings add to accumulating evidence that fast and slow spindles reflect distinct phenomena ^25–27,67^, and indicate that while these spindle types show distinct topographical patterns ^22–24^, their activity is modulated by the SO cycle even at sites where they are less prominent.

Second, coupling phase differed between sleep stages. Slow spindles occurred significantly later (i.e., 45° closer to the SO trough) in N2 than N3, while for fast spindles no important differences were found. Third, we observed subtle, but consistent regional differences in the phase of spindle expression (Fig. 6; Table 4). In particular, N3 spindles occurred ~25° later in the SO cycle in anterior vs. central regions, for both spindle classes. These differences were not explained by regional variability in SO amplitude, suggesting no direct link between the level of cortical polarization and spindle timing at the level of scalp EEG. Thus, while it is presently unclear what underlies these regional and stage differences, these factors are important to consider when the precise coupling phase is the focus of attention.

Beside systematic group-level variability, we found remarkably large individual differences in the preferred phase of spindle expression that, depending on condition, spanned up to half of the SO cycle, thereby attenuating group effects (e.g., non-significant group-level clustering of slow spindles: Fig. 4CD). Intriguingly, this variability was highly stable within subjects across nights (Fig. 8), and even observed within cortical regions (Fig. 9). Several features of SO ^45^ and spindle activity ^22,28,68^ show large yet reproducible individual differences related to underlying variability in anatomy ^29^. From this perspective, the fingerprint-like nature of SO-spindle coupling is perhaps not entirely surprising. But like regional variability, individual differences in coupling phase were unrelated to SO amplitude, leaving the cause of this heterogeneity unknown. However, previous reports correlating coupling phase to memory change rely on the implicit assumption that coupling phase is a state-like phenomenon in which individual differences are exclusively related to consolidation of recently encoded information. In contrast, our results indicate that any observed relation between coupling phase and behavior may reflect a general trait, rather than a task-induced phenomenon, similar to other trait-like relations between sleep physiology and cognitive ability ^69–72^. Consistent with this notion, we found no differences in the SO phase of maximal spindle activity between the baseline and learning nights.

Although overnight MST performance improvements were consistent with prior studies ^38,50–52^, we did not find any link between coupling phase and memory change, contrasting with several recent observations ^16,38,39^. However, the aforementioned studies are rather inconsistent, showing higher performance in either later ^16,38^ or earlier ^39^ coupling phases, following pharmacological spindle enhancement but minimally after placebo ^39^, and in patients but not controls ^38^. Hence, it remains to be seen whether, or under what conditions, coupling phase mediates memory consolidation.

SO densities varied considerably across the scalp (Fig. 2A, Table 2), resulting in both noisier within-subject coupling estimates and smaller sample sizes in regions with fewer SOs (Fig. 2D), potentially explaining the weaker effects in posterior regions. While we employed relatively lenient SO detection criteria, future work may examine whether further lowering amplitude thresholds, thereby increasing the number of detected SOs, would result in more robust posterior effects. Although topographical variability in SO density could be interpreted as variability in SO-spindle coupling, we assessed coupling for individually detected SOs to avoid this confounding influence. Thus, while the likelihood of observing SOs clearly varies across the scalp, whenever and wherever they are detected, SOs robustly organize the expression of local spindle activity.

SO-spindle coupling was determined separately at each of 58 electrodes, contrasting with studies assessing coupling on a limited number of channels ^30,38,73^, and with approaches where multi-channel spindle activity is related to the SO phase from a single (virtual) channel ^30,32,33^. The latter approach assumes that SOs are relatively global events that are synchronized across the scalp. While a minority of scalp SOs may indeed be global ^46^, our analyses of SO co-occurrence (≤400 ms) indicate that the average SO involves fewer than ten channels, consistent with invasive findings ^20^. Counting only SOs that co-occur with minimal phase shifts (≤100 ms; Fig. 2C, 3) resulted in a 40–45% reduction in channel involvement, further arguing against SOs as a uniform, zero phase-lag phenomenon. Such phase shifts are also consistent with evidence of SOs propagating across the cortex ^20,45,74^.

Local EEG dynamics were accentuated with the surface Laplacian ^58,59^. Although it might be argued that this approach suppresses true global SO and spindle activity, available evidence suggests that the Laplacian results in no loss of global information ^60,75^, while also providing improved estimates of oscillatory phase ^60^. More importantly, compelling evidence indicates that SOs and spindles have important local components, at the spatial scales of both invasive ^20,41–44^ and macroscopic EEG ^31,45,46^ (Fig. 1) recordings, strongly favoring an approach sensitive to such features. We speculate that the more global appearance of SOs in spatially unfiltered EEG recordings is at least partly due to volume conduction, ensuring that large-amplitude frontal SOs are detected at distant sites. Future studies may employ multi-resolution EEG approaches ^60,76^ to offer a fuller description of oscillatory sleep dynamics at different spatial scales.

We assessed coupling between SOs and continuous fluctuations in sigma power rather than individually detected sleep spindles. Because the spatiotemporal properties of SO-sigma coupling are relatively similar in the presence and absence of discrete spindles ^32^, the analysis of continuous sigma activity arguably offers a more comprehensive perspective on coupling dynamics. Nonetheless, our results are in line with reports based on discrete spindle detection ^16,30,32,38^, suggesting that this methodological choice does not pose a major concern.

Finally, we emphasize that examinations of preferred coupling phase only partially capture the dynamics of coupled sleep oscillations. Even when considering a single channel for a single subject, we have anecdotally observed that algorithmically detected spindles (1) reach their maximum amplitude in highly variable SO phases; (2) span a substantial portion or even the entire cycle of the SO waveform; or (3) may not be associated with any SO at all, most commonly during N2. Conversely, not all SOs are associated with discrete spindles. Given both our main findings and these additional complexities, we caution against the overly simplistic conceptualization that spindle activity is rigidly tied to a single SO phase. Still, spindle activity occurring in a narrow SO phase may induce plasticity more effectively ^40^, raising the possibility that the optimal phase could be different for different individuals and cortical regions. This suggestion may also be of practical relevance to closed-loop approaches targeting stimulus delivery at a specific SO phase ^5,8,77^.

In summary, we have identified systematic differences in the preferred SO phase of spindle expression as a function of spindle class, cortical region, sleep stage, and individual. While the causes and consequences of these many sources of variability remain to be determined, we suggest that locally coordinated oscillatory rhythms offer the sleeping brain a vast repertoire of building blocks to flexibly process, consolidate, and reorganize experiences encoded within and across brain structures, with important functional and clinical implications.

## Acknowledgments

This work was supported by grants from The Netherlands Organization for Scientific Research (NWO) to RC (446-14-009); National Institutes of Health to RS (MH048832), DSM (Manoach) (K24MH099421), RS and DSM (Manoach) (MH092638); George and Marie Vergottis Postdoctoral Fellowship to DSM (Mylonas); The Harvard Clinical and Translational Science Center (TR001102); and Stanley Center for Psychiatric Research at Broad Institute.

